# Enzymatic production of single molecule FISH and RNA capture probes

**DOI:** 10.1101/107599

**Authors:** Imre Gaspar, Frank Wippich, Anne Ephrussi

## Abstract

Arrays of singly-labelled short oligonucleotides that hybridize to a specific target revolutionized RNA biology, enabling quantitative, single molecule microscopy analysis and high efficiency RNA/RNP capture. Here, we describe a simple and efficient method that allows flexible functionalization of inexpensive DNA oligonucleotides by different fluorescent dyes or biotin using terminal deoxynucleotidyl transferase and custom-made functional group conjugated dideoxy-UTP. We show that 1) all steps of the oligonucleotide labelling – including conjugation, enzymatic synthesis and product purification – can be performed in a standard biology laboratory, 2) the process yields >90 %, often >95 % labeled product with minimal carry-over of impurities and 3) the oligonucleotides can be labeled with different dyes or biotin, allowing single molecule FISH or RNA affinity purification to be performed.

**Graphical abstract:** 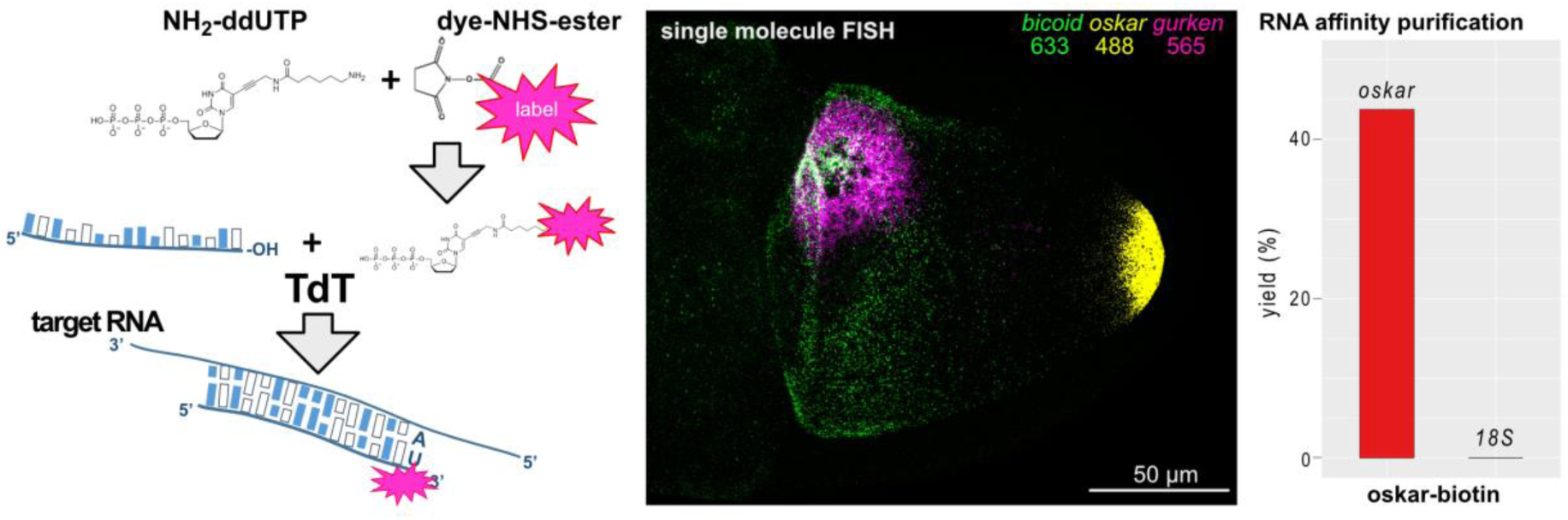

## Introduction

The ability to quantify biomolecules in space and time in living matter has introduced a qualitative change in many fields of biology allowing the building of accurate mathematical models that can test, validate and refine hypotheses about biological processes. Quantitative imaging of RNA, for instance, allows precise measurement of transcript copy number within cells, subcellular compartments, and ultimately in single ribonucleoprotein (RNP) complexes (1,2). Such studies have identified heterogeneity at all these levels: between cells (3), subcellular regions (4) and between individual RNPs. While some mRNPs contain only a single type and a single copy of an mRNA molecule, others may undergo dynamic homo- or heterotypic oligomerization, resulting in hundreds of copies of the mRNA within a single RNP particle (5). Although today a wide variety of RNA detection techniques with single molecule sensitivity exists (1), the pioneering techniques of single molecule fluorescent *in situ* hybridization (smFISH) were based on an array of short oligonucleotides carrying a well-defined number of labels, thus reducing variance of the detected signal while maintaining a high signal-to-noise ratio (6,7).

Such arrays typically consist of 24 to 96 different probes: 18-22 nucleotide long single stranded DNA (ssDNA) molecules complementary to non-overlapping segments of the target RNA, each carrying a single label (7). Due to the multiple probe molecules hybridizing to a single target, this design ensures a linearly amplified signal. As the vast majority (>90 %) of the probes carry only a single covalently coupled fluorescent molecule, heterogeneity of signal due to labelling is minimal in the case of smFISH probes, in contrast to conventional FISH probes. Furthermore, the relatively large number of probes ensures reliable detection of virtually all transcripts found within the specimen.

Due to the strict design principles, these probes are synthesized chemically and then purified by analytical means, such as HPLC or PAGE isolation (7). The coupling of the fluorophore takes place before purification and the resulting probe or probe set is rather static, such that there is no simple means to change to a spectrally different fluorophore, and it is often difficult or impossible to alter the composition of the oligonucleotides in the set without restarting the entire synthesis process.

An alternative to pre-coupled fluorophore probe sets are functionalized oligos that can be chemically coupled to a given molecule by the experimenter. The most commonly used functionalization is a primary amine modification at either end of the oligonucleotides, which enables simple coupling of dye-succinimidyl (NHS) -ester conjugates. In this way, aliquots of the same probe set can be labelled differently depending on the experimental requirements. However, such custom labelling puts the burden of purification – i.e. removing the unconjugated fraction of the oligos and the free dye molecules – on the experimenter. Also, the reactivity of NHS-esters decays rapidly as a function of storage time due to hydrolysis, which can result in unpredictable labelling and wasting of expensive reagents.

Here we present a simple, efficient and flexible alternative to smFISH and RNA affinity purification (RAP) probe production that can be carried out in any laboratory using basic equipment. This method relies on the template independent addition of dNTPs at the 3′ end of DNA by terminal deoxynucleotidyl transferase (TdT) (8). This enzyme could elongate 3′ ends indefinitely, however, the use of dideoxy nucleotides ensures that only a single labelled nucleotide is incorporated into each and almost every oligonucleotide molecule added to the reaction. Although the feasibility of such a direct labelling strategy was demonstrated previously for individual oligonucleotides (9,10), here we show that it is possible to label sets of conventional PCR oligos using custom-made dye/biotin conjugated ddUTPs synthesized in a standard biology laboratory. The synthesis results in >90 % (often >95 %) of singly labelled probe molecules, which - after a simple precipitation based purification – are suitable for quantitative imaging with single molecule sensitivity and for RAP. We describe the optimized conditions for incorporation of spectrally different Atto565-ddUTP and Atto633-ddUTP as well as biotin-ddUTP into ssDNA oligonucleotides.

## Results

### Incorporation of unpurified fluorophore-ddUTP conjugates to the 3′ of oligonucleotides by TdT

To make enzymatic oligo end-labelling a technique readily available for most laboratories, we identified simplicity and cost efficiency – as well as synthesis efficiency – as the main criteria for such an assay. Therefore, we first tested if TdT could incorporate unpurified Atto565-ddUTP, synthesized through conjugation of Atto565-NHS ester to NH_2_-ddUTP, into the 3′ end of single DNA oligonucleotides. We found that, given sufficient time (~7h), even relatively low amounts of the enzyme can yield near-qualitative labelling of both oskCD#16 and oskCD#6 oligos, as visualized upon denaturing 8 M Urea-PAGE (Figure 1A and B). The labelled fraction of the oligos yielded fluorescence in the red spectrum and, due to the addition of a nucleotide plus a bulky, positively charged dye, migrated more slowly during electrophoresis. This migratory retardation is large enough to distinguish incorporation of unlabeled NH_2_-ddUTP and dye-conjugated-ddUTP (Figure 1 and S1). As NHS-esters are known to lose reactivity rapidly in solution, we conjugated NH_2_-ddUTP with aliquots of Atto633 (2-fold molar excess) and Atto488 (10-fold molar excess) that were several (5-7) months old. As shown in Figure 1B, lane 4 and Figure 1C, lane 3, virtually all the oligonucleotide was labelled (i.e. the band corresponding to the unlabeled nucleic acid disappeared); however, in addition to a fluorescent, slow migrating band (Figure 1B, yellow arrowhead), a non-fluorescent product could be detected that reflects the incorporation of unlabeled NH_2_-ddUTP. Increased concentrations of DMSO (30 v/v %) – as a result of a large molar excess of the dye-NHS ester during conjugation – are detrimental to the synthesis (Figure 1C, lane 2), hence the reaction must be diluted to bring the DMSO concentration below 6 v/v % (Figure 1C, lane 3). The appearance of a middle band is an indicator of an incomplete conjugation reaction in the case of positively charged dyes (e.g. Atto488, Atto565 or Atto633). Unfortunately, this does not apply to negatively charged dyes, such as Atto495 or small, neutral compounds, such as biotin (Figure 1B and D) or BDP-FL (Figure S1)

**Figure 1:**
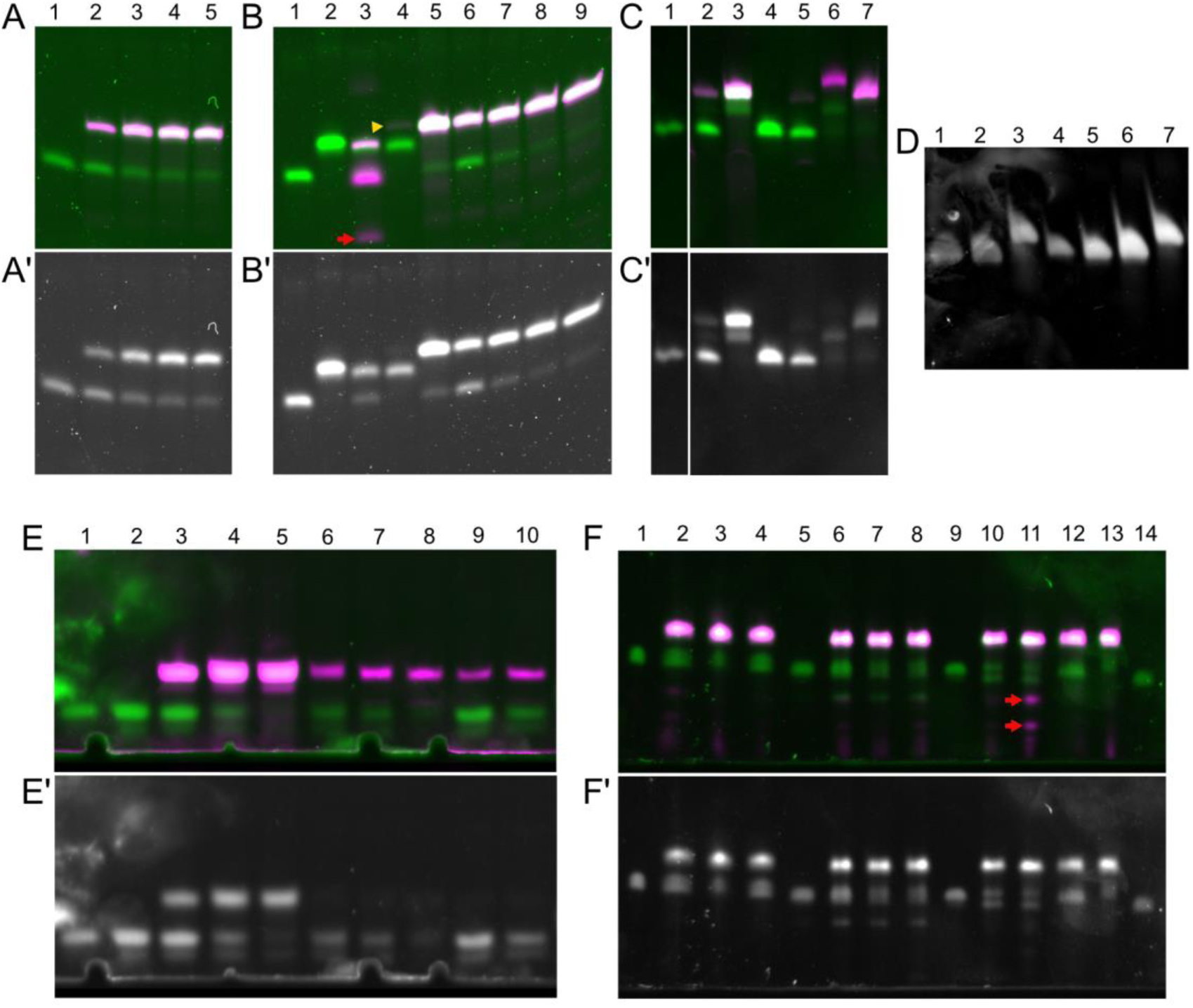
3′ incorporation of labelled ddUTP by TdT. A, A’ – oskCD#16 probe (1) labelled with Atto565-ddUTP (4x molar excess) for 1 h (2), 3 h (3), 5 h (4) and 7 h (5). Dye fluorescence is shown in magenta, SYBR GOLD staining is shown in green (A) and in gray (A’). 96 pmol oligo/lane. B, B’ - oskCD#6 probe (1) labelled with biotin-ddUTP (3-fold molar excess) (2), Atto495-ddUTP (4-fold molar excess) (3), Atto633-ddUTP (4-fold molar excess) (4) and Atto565-ddUTP (4-fold molar excess) (5) overnight (16h). Red arrow (3) indicates carry-over of free Atto495-ddUTP. Yellow arrowhead points to a minuscule amount of Atto633 labeled, fluorescent product in lane 4. Time course labelling of oskCD#6 probe with Atto565-ddUTP (4-fold molar excess) for 1 h (6), 3 h (7), 5 h (8) and 7 h (9). 96 pmol oligo/lane. C, C’ - oskCD#6 probe (1) labelled with Atto488-ddUTP (4-fold molar excess) in the presence of 30% (2) and 6% DMSO (3). oskCD#8 probe labelled directly with AttoRho14-ddUTP (4) and Atto725-ddUTP (5) (4-fold molar excess) or with unconjugated NH2-ddUTP reacted to AttoRho14-NHS ester (6) or Atto725-NHS ester (7) subsequently. 96 pmol oligo/lane. D - osk19nt-9x probe mixture (1 – 12 pmol, 2 – 24 pmol) labeled with biotin-ddUTP (3-fold molar excess) (3 – 24 pmol). osk20nt-15x probe mixture (4 – 6 pmol, 5 – 12 pmol, 6 – 24 pmol) labeled with biotin-ddUTP (3-fold molar excess) (7 – 24 pmol). E, E’ - osk20nt-15x probe mixture (1 – 6 pmol, 2 – 9 pmol) labelled with Atto565-ddUTP (3 – 1.5-fold, 4 - 2.5-fold, 5 – 5-fold molar excess) or with Atto633-ddUTP (6 – 1.5-fold, 7 – 2-fold, 8-10 – 2.5-fold molar excess). Lane 9 and 10 shows results of labelling performed with reduced amounts of TdT enzyme (9 – 0.2-fold, 10 – 0.5-fold of standard TdT amount). 15 pmol oligo/ lane (3-10). F,F’ – Atto565-ddUTP labelling (5-fold excess) of probe mixtures. gfp20nt-7x probe mixture (1) labelled using 1-fold (2,3) or 2-fold (4) the standard TdT amount. 18S20nt-31x probe mixture (5) labelled using 1-fold (6,7) or 2-fold (8) the standard TdT amount. osk20nt-15x probe mixture (9) labelled using 1-fold (10) or 2-fold (11) of standard TdT amount. osk19nt-9x probe mixture (14) labelled using 1-fold (12) or 2-fold (13) the standard TdT amount. Lanes 3 and 7 show results of re-labelling of probe mixtures shown in lanes 2 and 6, respectively. 3 pmol oligo/lane (1, 5, 9 and 14) and 15 pmol oligo/lane (2-4, 6-8 and 10-13). Red arrow (11) indicates carry-over of free Atto565-ddUTP.

Other dyes, such as AttoRho14 and Atto725, on the other hand, were readily conjugated to primary amines, but the resulting dye-ddUTP product was not incorporated during the synthesis (Figure 1C, lanes 4 and 5). Nevertheless, it was possible to use the NHS-ester of these dyes to conjugate NH2-ddUTP labelled oligos (Figure 1C, lanes 6 and 7).

### Enzymatic production of smFISH and RNA capture probes

For efficient and relatively simple synthesis of smFISH and RNA capture probe sets, the assay should yield >90 %, ideally >95 % of labelled oligonucleotides (as offered by vendors of smFISH probes) with minimal contamination of the free dye-ddUTP conjugate. Also, the assay should be capable of labelling not only single oligos, but also pools of different oligonucleotides. To fulfill these criteria, we optimized the labelling reactions by varying some of the experimental conditions, for instance using different probe sets containing oligos of the same length (osk-19nt-9x, osk-20nt-15x, gfp-20nt-7x, 18S-20nt-31x). We found that efficient incorporation of the different ddUTP conjugates required slightly different conditions: a 3-fold excess of biotin-ddUTP and standard amounts of TdT were sufficient to label virtually all molecules of the probe sets (Figure 1D). In the case of Atto633-ddUTP, we needed as little as a 2.5-fold excess (although, as a precaution, in our protocols we use 3-fold excess of this nucleotide) and a standard amount of TdT to obtain near qualitative labelling (Figure 1E). However, sub-standard amounts of the enzyme negatively impacted the synthesis (Figure 1E, lanes 9-10). In the case of Atto565-ddUTP, we used a 5-fold molar excess to obtain similar results (Figure 1E). Although a standard amount of the enzyme worked sufficiently in case of the osk-20nt-15x probe set, efficient labeling of other probe sets (e.g. 18S-20nt-31x or osk-19nt-9x) required twice as much enzyme as the standard (Figure 1F). In case of Atto633-ddUTP, 3-fold molar excess of the ddUTP conjugate and standard amounts of TdT worked for all other probe sets (data not shown). Efficient labelling of the gfp-20nt-7x probe set with Atto565-ddUTP turned out to be especially difficult, probably due to the intra- and intermolecular interactions of the individual oligonucleotides, which had high melting temperature (T_m_) as a consequence of the GC richness of the *gfp* coding sequence. However, we found that subjecting the labelled oligo mixture to a second round of labelling (using standard amounts of TdT in both synthesis steps) could yield >90 % labelled gfp-20nt-7x product (Figure 1F, lanes 2, 3 and 6, 7).

Although there are different ways to purify the oligos after the synthesis (e.g. gel filtration, HPLC or PAGE purification), we found that for most of the dye-ddUTP conjugates - with the exception of Atto 495 (Figure 1B, red arrow), Abberior 470 SX, Atto488 and AlexaFluor488 (Figure S1, red arrow) - a simple, linear acrylamide facilitated ethanol precipitation and subsequent washes usually resulted in a clean product with little or no visible trace of the free dye or the dye-ddUTP conjugate (Figure 1F, lane 11, red arrows). Such purification further simplifies the handling of these oligo mixtures and, starting with 1000 pmol of unlabeled oligo, the typical loss is 22.0 ± 12.9% (mean ± SD, N = 33). Another implication of the low carry-over of fluorescent impurities is that the degree-of-labelling (DOL, i.e. the fraction of labelled oligos) determined by spectroscopy of the reconstituted oligo mixture is expected to correlate well with that measured by densitometry on PAGE. To test this hypothesis, we plotted the spectroscopic DOL as a function of densitometric DOL of the dye-labelled oligo mixtures and fitted the plots with a linear regression model (Figure 2A). We found a great fit between our data and the linear model (R^2^>0.98) with a slope slightly above 1 for both Atto565 and Atto633 labelling, and in the vast majority cases the ratio of the two DOL values was within 10 % of the mean (Figure 2B). This indicates that spectroscopy provides a good – although slightly overestimating - measure of the DOL of these enzymatically produced smFISH probes.

**Figure 2:**
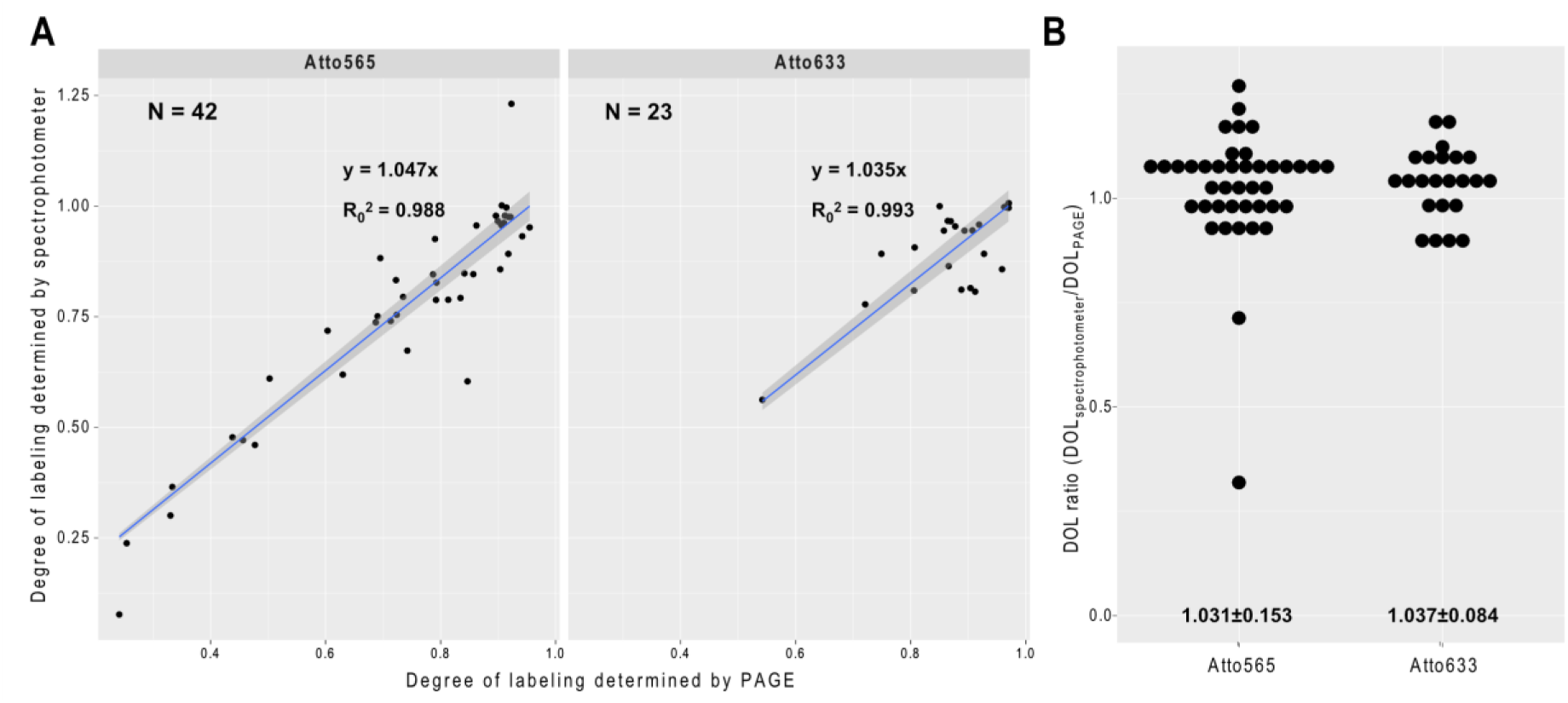
Determining degree of labelling. A – Degree-of-labelling of isomeric probe mixtures (osk19nt-7x, osk20nt-15x, gfp-19nt-11x, gfp-20nt-7x and 18S-20nt-31x) determined by PAGE densitometry and by spectrophotometry using Atto565- or Atto633-ddUTP. Correlation of the two analytical modalities was tested, slope and goodness-of-fit, as well as the number of measurements are indicated in the graphs. The intercept of the regression model was set to zero. B – Ratio of spectroscopically and densitometricly determined degree-of-labelling. Mean ± SD are indicated below the graphs.

### Assaying performance of enzymatically produced smFISH probes

smFISH probes are expected to detect even a single copy of their target mRNA with high sensitivity (low level of false negatives) and with high specificity (low level of false positives). To test the performance of the labelled probe sets, we chose the developing *Drosophila* egg-chamber as a model system: this tissue is the unit of *Drosophila* oogenesis that with time produces oocytes of a several hundred micron scale, rendering this model challenging for microscopy. On the other hand, a single egg-chamber is composed of two tissue-types: germline (oocyte and the nurse cells) and soma (the follicular epithelium that surrounds the germline). As these cells differ in origin and function, and each has a plethora of cell-type specific transcripts, they constitute ideal internal negative controls for each other in the smFISH assay.

To test the single molecule sensitivity of our probe sets, we labelled the germ-line specific *oskar* mRNA with two different fluorophores coupled to two probe sets (osk18x-Atto633 and osk17x-Atto565, the probes are arranged in an alternating setup, Table S1). As shown in Figure 3A, there was a high degree of overlap between the two colors in the germ-line, but hardly any puncta were observed in the follicle cell layer with either of the fluorophores. Similarly, another germ-line specific mRNA, *nanos* was detected almost exclusively in the nurse cells, however no apparent co-localization was observed between the nos18x-Atto633 and osk17x-Atto565, consistent with previous observations (5) indicating specificity of these probe sets (Figure 3B).

**Figure 3.**
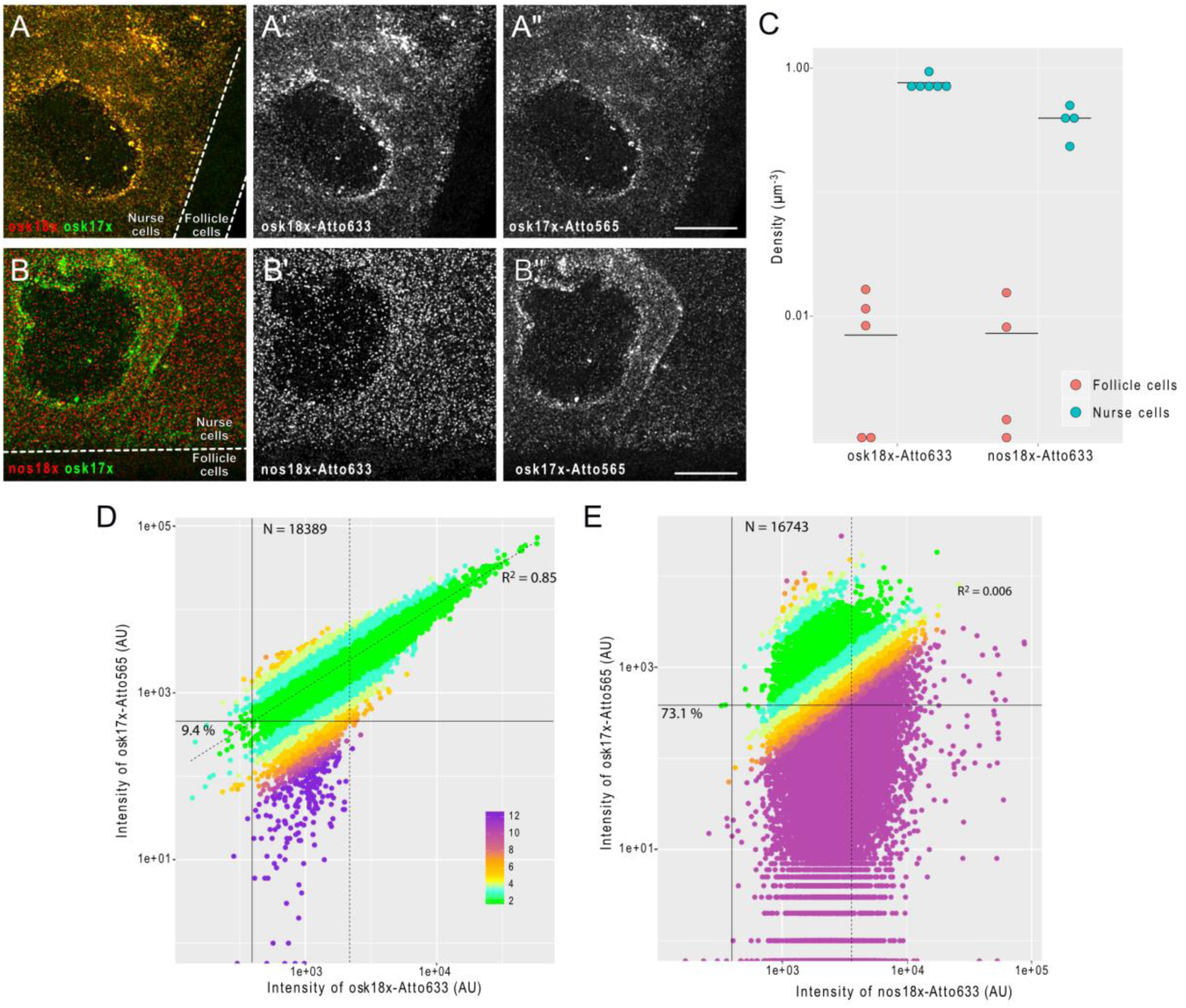
smFISH carried out using 3′ end labelled probe mixtures. A-A” – smFISH of *oskar* mRNA in a developing *Drosophila* egg-chamber using osk18x-Atto633 (red, A’) and osk17x-Atto565 (green, A”) probe sets. The two probes sets label *oskar* mRNA in an alternating fashion. B-B” - smFISH of *oskar* and *nanos* mRNAs in a developing *Drosophila* egg-chamber using nos18x-Atto633 (red, A’) and osk17x-Atto565 (green, A”) probe sets. A-B” - The germline, which expresses *oskar* and *nanos* mRNAs (nurse cells) and the soma (follicle cells) are indicated. Scale bars are 10 µm. C – Density of detected smFISH signal in the mRNA expressing (cyan, nurse cells) and non-expressing (red, follicle cells) compartments of the egg-chambers. Horizontal lines indicate the mean value of observations. D, E – Intensity of osk17x-Atto565 probe sets (target channel) as a function of Atto633 signal intensity (reference channel) of smFISH objects detected by the fluorescence of osk18x-Atto633 (D) or nos18x-Atto633 (E) probe sets (reference channel). Percentage values represent the fraction of single mRNA containing smFISH objects that would have failed to be detected in the target channel (Atto 565) since their corresponding signal intensity falls below the estimated target channel detection threshold (solid horizontal line). This detection threshold corresponds to the product of the reference channel detection threshold (solid vertical line, 0.1^th^ percentile of the Atto633 signal intensity distribution) and the slope of the fitted line (dashed line. The line was fitted to the non-single mRNA representing fraction of the population; goodness-of-fit indicated). smFISH objects with signal intensity lower than µ_1_+2σ_1_ of the smallest fitted Gaussian function (Figure S2) were considered to represent a single mRNA molecule (dashed vertical line). Numbers of these single copy mRNA objects are indicated in the graphs. Colors represent the relative fold difference of the observed and expected target channel (Atto565) signals (see panel D for key). The expected target channel (Atto565) signal is calculated as the product of the regression slope and the reference channel (Atto633) fluorescence.

We subsequently analyzed automatically detected smFISH objects using one of the colors as the reference the other as the target channel. By comparing the object densities, we found 20-100-fold more objects in the transcript expressing germ-line relative to the non-expressing follicle cell layer (Figure 3C and Figure S3B), indicating that the false positive detection rate (FPDR) of these probe sets is around 5 - 1 %. Next, we assayed the single molecule co-detection rate of the two differently labelled probe sets. As in the nurse cells *oskar* mRNA was shown to be present in RNPs containing single and two copies of the mRNA (5), we first fitted the signal intensity distribution of the reference channel with Gaussian functions as described in (5)(Figure S2). We considered that objects with a fluorescence intensity lower than the µ+2σ of the first fitted Gaussian function contained a single mRNA molecule (Figure 3D and E and Figure S2, dashed vertical lines). Within this population of objects, we estimated a detection threshold of the target channel (see legend of Figure 3D and E). We found that ~ 91 % of objects detected by osk18x-Atto633 would be co-detected by osk17x-Atto565 - ~86 % co-detection rate in the complementary analysis (Figure S3C). These values are in good agreement with what has been previously observed using chemically synthesized smFISH probes (7,11,12). On the other hand, only 27 % of nos18x-Atto633 positive objects showed detectable osk17x-Atto565 fluorescence, and there was no linear relation between the signal intensities of these two probes (R^2^ = 0.006).

To assess the performance of our gfp probe set, we performed similar co-detection analysis on egg-chambers that expressed *oskar-EGFP* mRNA in addition to the endogenous *oskar* mRNA (Figure S3A). Despite the relatively high FPDR of our gfp23x-Atto633 probe set (~10 %, Figure S3B) – probably due to the considerably higher T_m_ of the gfp probes than the osk probes – we found that ~78 % of gfp23x-Atto633 positive objects were co-detected by osk42x-Atto565 (Figure S3D).

Biotinylated DNA probes have been used previously to isolate RNA from complex samples by in solution hybridization (13-16). Therefore, we wanted to assess the performance of our biotinylated DNA probe set for osk (osk24x-biotin) to purify endogenous *oskar* mRNA from *Drosophila* ovarian lysates. Compared to a single 400 nucleotide long *in vitro* synthesized antisense RNA probe that has incorporated multiple biotinylated UTPs (approx. 11 per RNA) at various positions, equimolar amounts of osk24x-biotin probes were capable of capturing approximately 10-fold more endogenous *oskar* mRNA (Figure 4). Furthermore, we found that osk24x-biotin probes selectively captured the desired mRNA with very little cross-contamination, as judged by the amounts of co-purified 18S ribosomal RNA. Thus, self-made biotinylated DNA probes are cost efficient tools that can be used for biochemical purification of endogenous mRNAs for subsequent analysis.

**Figure 4.**
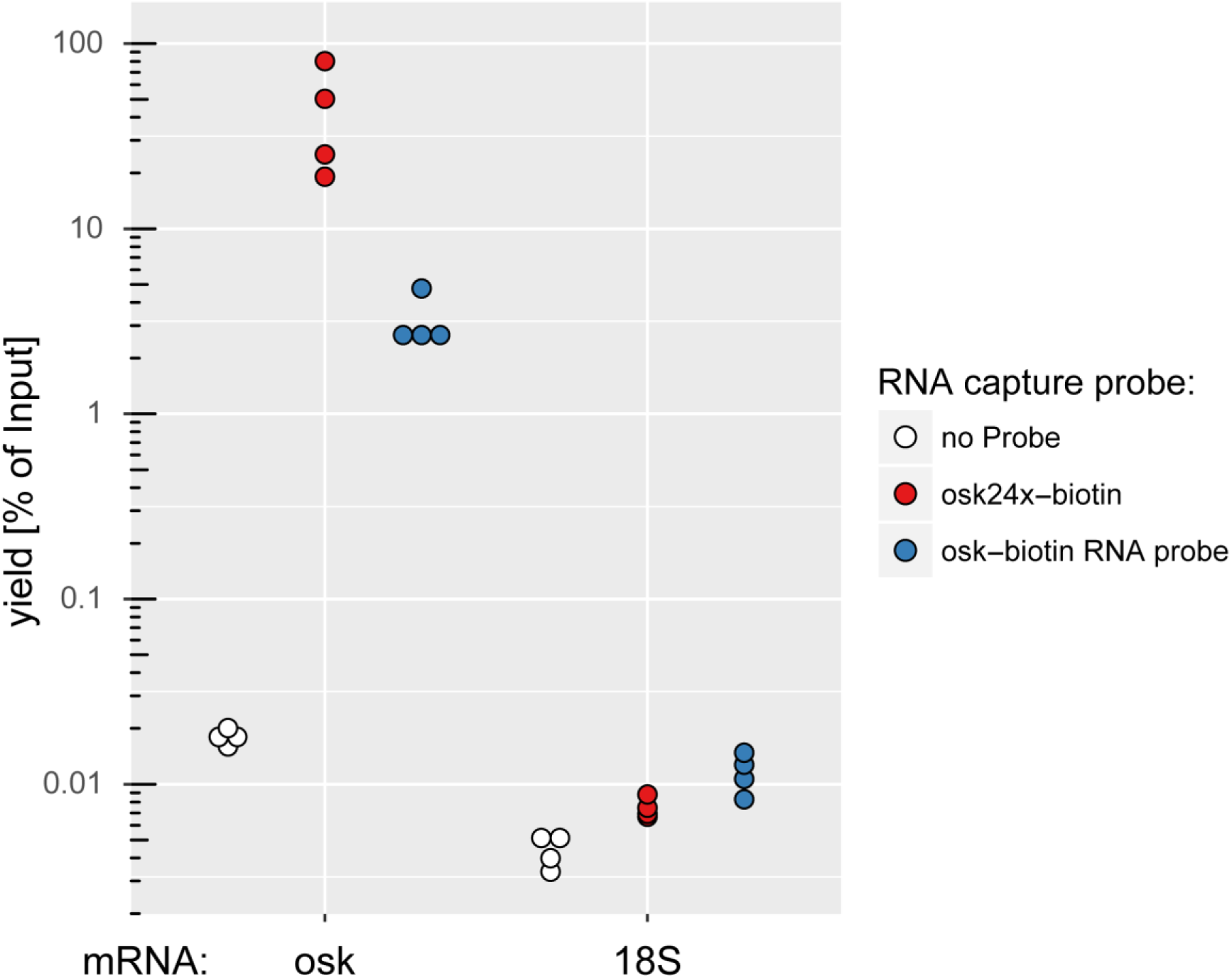
mRNA affinity purification using biotinylated DNA vs. RNA probes. A biotinylated DNA probe set (osk24x-biotin) captured *oskar* mRNA more efficiently than a single, 400 nucleotide long, internally biotinylated RNA probe (osk-biotin RNA probe), whereas the a unrelated RNA (18S ribosomal RNA) was only marginally captured.

## Discussion

In summary, the method we have presented here is capable of converting an array of inexpensive PCR oligos complementary to a given RNA target into functional smFISH or RAP probes adapted to the experimenter’s specific needs. Although we focused on the optimization of Atto565 and Atto633 labelling, as shown in Figure 1 and Figure S1, quite a number of the different dyes we tested are suitable for use with this method. Also, as exemplified by the different combinations of *oskar* probes we used in this and a previous (17) study, the method offers unprecedented flexibility in choosing and manipulating the composition of the generated probe sets. Importantly, as opposed to labelling primary amine containing oligonucleotides, here we convert the volatile dye-NHS ester into a stable dye/biotin-ddUTP conjugate that can be thawed/frozen multiple times and stored without perceptible loss of activity (our oldest aliquot in use is an over one year old Atto633-ddUTP conjugate).

Another important aspect of this method is the associated specific cost of a labelling reaction is low: a typical 1000 pmol scale synthesis of a set of 30 oligos usually yields probes sufficient for ~ 100 smFISH experiments (using 2.5 nM/probe final concentration in a 100 µL hybridization volume). The costs of the initial set of 30 ssDNA oligos (typical delivery amount is ~ 30 × 20 000 pmol) is usually a two-digit figure (in USD). The expensive components of the labelling reaction are sufficient for ~40-80 (TdT, 500 U), ~100 (1 µmol Atto565-NHS ester) and ~200 (1 µmol ddUTP) of such 1000 pmol scale reactions, and thus cost below 10 USD including production and purification.

Taken together, as the entire synthesis and purification takes ~ 24 h and requires no special equipment or prior expertise, we consider this method a good alternative to commercially available smFISH probe sets.

## Methods and Materials

### Synthesis of labelled ddUTP

NH_2_-ddUTP (Lumiprobe, stock concentration 10 mM) was combined with two-fold molar excess of dye-NHS-ester in the presence of 0.1 M NaHCO_3_, pH 8.3 and the reaction mixture was incubated for 2-3 hours at room temperature isolated from light. Subsequently, 1 M Tris HCl, pH 7.4 was added to 10 mM final concentration in order to quench any unreacted NHS-ester groups. The reaction mixture was adjusted with nuclease-free H2O to obtain 2-5 mM final concentration of the labelled-ddUTP. The following dye-NHS-esters were used: Atto425-NHS, Atto465-NHS, Atto488-NHS, Atto565-NHS, Atto633-NHS, AttoRho14-NHS, and Atto725-NHS were purchased from Atto-tec GmbH, Abberior 470SX-NHSl Abberior RED-NHS and Abberior 635P-NHS esters were a gift of Abberior GmbH; BDP-FL-NHS-ester was obtained from Lumiprobe. The NHS-esters were reconstituted with anhydrous DMSO to 20-40 mM concentration in a chamber filled with Blue Silica desiccant to prevent moisture contamination of the NHS-ester. AlexaFluor488-NHS-ester was obtained from LifeTechnologies Inc. and was reconstituted to 10 mM concentration. Biotin-NHS-ester was obtained from SigmaAldrich GmbH and reconstituted to 20-40 mM final concentration. Typically, the totality of the reconstituted NHS-ester was immediately put in reaction with NH_2_-ddUTP. In the initial experiments, several month old aliquots of Atto488-NHS and Atto633-NHS esters were used, reconstituted to 2 mM. Depending on the initial concentration of the reconstituted dye, the final DMSO content of the labelled-ddUTP was between 25 - 80 %.

### Enzymatic production of labelled oligonucleotides

Non-overlapping arrays of 18-22mer DNA oligos complementary to *oskar*, *nanos*, *gfp* and 18S RNA targets were manually selected (Table S1). The selection criteria included 45-60 % GC content, similar melting temperature, and the oligos were designed such that in most cases the 3′ incorporated ddU was also part of the hybrid. Adjacent oligos were separated by at least two nucleotides. Desalted or reverse phase cartridge purified PCR oligos were obtained from SigmaAldrich GmbH and reconstituted to 240-250 µM with nuclease-free H_2_O. 500-3000 pmol of individual or pooled oligonucleotides were mixed with 1.5-5-fold molar excess of the labelled ddUTPs (as indicated in the Results and in the Figure legends) in 1x TdT buffer. The mixture was completed by the addition of TdT enzyme (0.006 U/pmol ssDNA, standard amount) and was incubated at 37 °C typically overnight in a PCR thermocycler with hot-lid on. The provided Excel sheet (Table S2) provides the precise composition of the reaction mixture for the optimized Atto565, Atto633, and biotin labelling.

### Purification of labelled oligonucleotides

Upon completion of the enzymatic reaction,, the reaction mixtures were supplemented with Na-acetate, pH 5.5 (300 mM final concentration), 1.5-5.0 µg linear acrylamide (Thermo Fisher Scientific) depending on the synthesis scale (e.g. 2.5 µg for a 1000 pmol, or 5 µg for a 3000 pmol scale synthesis), and diluted to 200 µL with nuclease-free H_2_O. This mix was transferred into a conventional 1.5 mL Eppendorf tube (DNA low binding tubes appear bind the non-incorporated free dyes) and 800 µL – 20 °C 100 % ethanol was added and mixed well. Precipitation was facilitated by incubating the mixture at – 80 °C for 15-20 minutes. Precipitated oligos were pelleted by centrifuging with 16000x g at 4 °C for 20 minutes. The pellet was washed in 1 mL 4 °C 80 % ethanol by vortexing and pelleting with 16000x g at 4 °C for 20 minutes. After the first wash, the pellet was transferred into a clean 1.5 mL Eppendorf tube and the washing was repeated two more times. Subsequently, residual traces of ethanol were removed and the pellet was air-dried before reconstitution with nuclease-free H_2_O.

### Analysis of the labelled oligonucleotides

Degree-of-labelling (*DOL*), concentration (*c*) and recovery efficiency (*recovery%*) of the labelled oligonucleotides was assessed by UV-Vis spectroscopy using a NanoDrop spectrophotometer. Absorbance at 260 nm (OD_260_) and at the dye absorption maximum (OD_dye_) were measured and the following formulae were used to estimate properties of the labelled oligonucleotide:

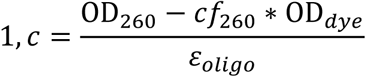

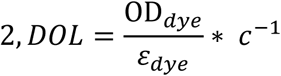

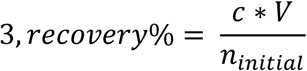

*ε*_*dye*_ and *cf*_260_ are the molar extinction coefficient and the relative absorbance at 260 nm of the dye, respectively. *ε*_*oligo*_ is calculated by taking the average of the extinction coefficient of oligonucleotides in the oligo mixture and adding 9000 mol^-1^cm^-1^ corresponding to the incorporated ddU. *V* is the reconstituted volume of the labelled oligo, *n*_*initial*_ is the starting amount of unlabeled oligo.

To determine the degree of incorporation of biotin we used the HABA/Avidin assay (SigmaAldrich) that is based on the displacement of HABA from avidin by biotin, resulting in a loss of color that can be quantified by measuring the absorbance at 500 nm. In the case of all our biotinylated probe sets, we measured close to equimolar amounts of biotin and oligonucleotides present in the samples (DOL ~ 1).

Denaturing PAGE analysis was performed on labelled single oligonucleotides or pooled oligonucleotides of the same length. 15-96 pmol of labelled product were denaturated by boiling in presence of 4 M urea and 1x DNA loading dye for 10 minutes and were immediately loaded to 15 % polyacrylamide gel containing 1x TBE and 8M Urea. The gel was pre-run for at least 30 minutes at 30V/cm and, once loaded, the same voltage was used to separate the denatured oligonucleotides using the unlabeled oligo as size marker. Gels were imaged with a BioRad ChemiDoc gel documentation system using 254 nm excitation and long-pass visible light filter to detect dye fluorescence. Subsequently, gels were stained with 2X SYBR GOLD in 1x TBE for 10 minutes and imaged with a Peqlab gel documentation system using 254 nm excitation and a band-pass visible light filter. This allowed visualization of the unlabeled fraction of the oligo in addition to the labelled product. Due to the band-pass filter used and the possibly highly efficient energy transfer between SYBR GOLD and the far red dyes, the Atto633 and Abberior RED labelled products are not visible in the stained gels (Figure 1E’ and Figure S1). The two images (before and after staining) were aligned in ImageJ using the gel boundaries as transformation landmarks. To determine DOL by PAGE, the amount of unlabeled oligo (*n*_*unlabelled*_) was determined by comparing the fluorescence of the corresponding band to a series unlabeled oligo of known quantity (typically 3, 6 and 9 pmol). DOL was calculated as follows: *1 - n*_*unlabelled/nloaded*_

### smFISH and image analysis

The following probe sets were used during smFISH: osk17x-Atto565 (DOL = 0.93), osk18x-Atto633 (DOL=1.07), nos18x-Atto633 (DOL=1.07), osk42x-Atto565 (DOL=0.96), gfp23x-Atto633 (DOL=0.94). Single molecule FISH was performed similarly as described in (17) using ovaries of *w^1118^* and *oskar-EGFP* expressing (18) female flies. Briefly, ovaries were dissected into 2 v/v % PFA, 0.05 v/v % Triton-X-100 in PBS (pH 7.4) and were fixed for 20 minutes on an orbital shaker. The fixative was removed and the ovaries were washed twice in PBT (PBS + 0.1 v/v % Triton-X-100, pH 7.4) for five minutes. *w^1118^* samples were treated with 2 µg/mL proteinase K in PBT for 5 minutes and then were subjected to 95 °C in PBS + 0.05 v/v % SDS for five minutes. Specimens were cooled by adding 2x volume of room temperature PBT. Proteinase K/heat treatment was omitted in the case of *oskar-EGFP* expressing samples so as to preserve GFP fluorescence. Ovaries were prehybridized in 200 µL 2xHYBEC (300 mM NaCl, 30 mM Sodium citrate pH 7.0, 15 v/v % ethylene carbonate, 1mM EDTA, 50 µg/mL heparine, 100 µg/mL salmon sperm DNA, 1 v/v % Triton-X-100) for 10 minutes at 42 °C. 50 µL of pre-warmed probe mixture (12.5-25 nM/individual oligonucleotide) was added to the prehybridization mixture and hybridization was allowed to proceed for 2 hours at 42 °C. Free probe molecules were washed out of the specimen by a series of washes: 0.5 mL pre-warmed 2xHYBEC, 1 mL pre-warmed 2xHYBEC:PBT 1:1 mixture, 1 mL pre-warmed PBT for 10 minute at 42 C and finally 1 mL pre-warmed PBT allowed to cool down to room temperature. Ovaries were mounted in 80 v/v % 2,2-thiodiethanol in PBS.

Stacks of images were acquired on a Leica TCS SP8 confocal microscope using a 63x 1.4 NA oil immersion objective and were restored by deconvolution in Huygens Essential. Deconvolved images were analyzed in ImageJ using a custom-made particle detection and tracking algorithm (17,19). Briefly, the algorithm finds in each slice of the reference channel the local maxima that represents the upper few percentile of the signal distribution. These 2D objects are then connected along the z-axis based on their center-positions to create 3D objects. Signal intensities of both the reference and target channels of all 3D objects with a minimum of 3 slices depth (smFISH object) found within the nurse cell compartment were recorded and were subject of statistical analyses in R (as described in the legend of Figure 3). To determine FPDR, smFISH object density in a manually selected regions in the follicle cells was compared to the smFISH object density in the nurse cells in randomly selected regions of comparable volumes.

### RNA Capture Method

Ovaries from well-fed *Drosophila melanogaster* (Oregon R) were resuspended in 3x volume of lysis buffer (50 mM Tris-HCl pH 7.0, 10 mM EDTA, 1 v/v % SDS, supplemented with fresh PMSF 1 mM, cOmplete^®^ mini EDTA-free protease inhibitor (Roche) and RiboLock RNAse Inhibitor (Thermo)) and mechanically homogenized. The lysates were cleared by centrifugation (5 minutes at 140x g). The supernatant was further diluted with 2 volumes (2:1) hybridization buffer (750 mM NaCl, 1 v/v % SDS, 50 mM Tris-HCl pH 7.0, 1 mM EDTA, 15 v/v % ethylene carbonate (Sigma), fresh 1mM PMSF, cOmplete^®^ mini EDTA-free protease inhibitor (Roche) and RiboLock RNAse Inhibitor (Thermo)), precleared with Pierce^®^ Avidin Agarose (Thermo) for 30 minutes and cleared by centrifugation (5 minutes at 140x g). The precleared lysates were supplemented with 0.25 µg of biotinylated probes (osk24x-biotin) or of a 400 nucleotide long biotinylated RNA complementary to the 3′ of the *oskar* cds (19) per mL ovaries and incubated at 37 °C for 2 hours with constant rotating. RNA-probe complexes were collected by the addition of magnetic Streptavidin beads (Dynabeads^®^ MyOne™ C1, Thermo) for 1 hour at 37 °C. The beads were washed 3 times for 5 minutes at 37°C with low salt wash buffer (300 mM NaCl, 30 mM Sodium citrate pH 7.0, 0.5 v/v % SDS, fresh PMSF and cOmplete^®^ mini EDTA-free protease inhibitor (Roche)), 2 × high salt wash buffer (750 mM NaCl, 30 mM Sodium citrate pH 7.0, 0.5 v/v % SDS, fresh PMSF and cOmplete^®^ mini EDTA-free protease inhibitor (Roche)) and 2 × low salt wash buffer. The RNA was eluted in TE buffer (10mM Tris-HCl pH 7.0, 1 mM EDTA) at 95°C for 5 minutes and extracted with Quick-RNA™ MicroPrep Kit (Zymo Research). cDNA was synthesized using SuperScript™ III First-Strand Synthesis SuperMix (Thermo) with random hexamer. Real-time PCR analysis was carried out using SYRB^®^ Green PCR Master Mix (Applied Biosystems) on a StepOnePlus™ Real-Time PCR System (Applied Biosystems) with primers against *oskar* (forward: 5′- CAGACTCTTCTCGTCCACTCAG-3′ and reverse: 5′-CGTGCAGTGGAAATGGATTGC-3′) or 18S rRNA (forward: 5′- CGGAGAGGGAGCCTGAGAA-3′, reverse: 5′- AGCTGGGAGTGGGTAATTTACG-3′. The results were compared to a dilution series of input samples to calculate the yield of the capturing procedure.

### Statistical analyses

All statistical analyses (indicated in the Results and in the Figure legends) were carried out in R (20) using RStudio (www.rstudio.com). All graphs were plotted by the ggplot2 library in R (21).

## Acknowledgements

### Author contributions

I. G. conceived and optimized the enzymatic, smFISH and RNA capture probe production method and performed the smFISH analyses. F. W. synthesized the RNA capture probes and performed the RNA affinity purification. I. G., F. W. and A. E. wrote the manuscript.

### Acknowledgment

We thank the EMBL Advanced Light Microscopy Facility and Leica for providing cutting-edge microscopy, and the EMBL Genomics Core Facility for their help with qRT-PCR. This work was funded by the EMBL. F.W. was supported by a postdoctoral fellowship from the EMBL Interdisciplinary Postdoc Program (EIPOD) under Marie Curie COFUND actions.

### Conflict of Interests

The authors declare that they have no conflict of interest.

